# Structural basis of substrate specificity of *Helix pomatia* AMP deaminase and a chimeric ADGF adenosine deaminase

**DOI:** 10.1101/2025.03.26.645602

**Authors:** Gundeep Kaur, John R. Horton, George Tzertzinis, Jujun Zhou, Ira Schildkraut, Xiaodong Cheng

**Author notes:** Email: GK, JRH, GT, JZ.

## Abstract

HPAMPD, an enzyme enriched in the foot muscle of the mollusk *Helix pomatia*, exhibits deaminase activity on adenosine-5’-monophosphate (AMP). HPAMPD is the first member of the <u>a</u>denosine <u>d</u>eaminase-related growth factor (ADGF) family to prefer the nucleotide, AMP, over the nucleoside, adenosine. To investigate the substrate selectivity of HPAMPD, we determined its structure in the apo form and in complex with the adenosine analogs pentostatin (2’- deoxycoformycin) and pentostatin-5’-monophosphate. Structurally, HPAMPD adopts a fold similar to human ADA2, an ADGF family member. HPAMPD has acquired the ability to interact with the 5’-monophosphate group of AMP through polar and charged residues located in three key structural elements: (1) the loop immediately following strand β1, (2) the loop between helices αH and αI, and (3) the end of strand β5 and its adjacent loop. We engineered a chimeric deaminase by integrating these elements from HPAMPD into another related mollusk nucleoside adenosine deaminase, the *A. californica* ADGF. The chimeric enzyme efficiently deaminates AMP, demonstrating a gained substrate specificity, while retaining the adenosine deamination activity of *Aplysia* ADGF. The phosphate-binding feature of HPAMPD is a hallmark of nucleotide deaminases, conserved among AMP and N6-methyl-AMP (6mAMP) deaminases. We discuss the human adenosine deaminases each with distinct substrate specificities for the nucleoside, the nucleotide (AMP), and the methylated form, 6mAMP.

**Graphical Abstract:** 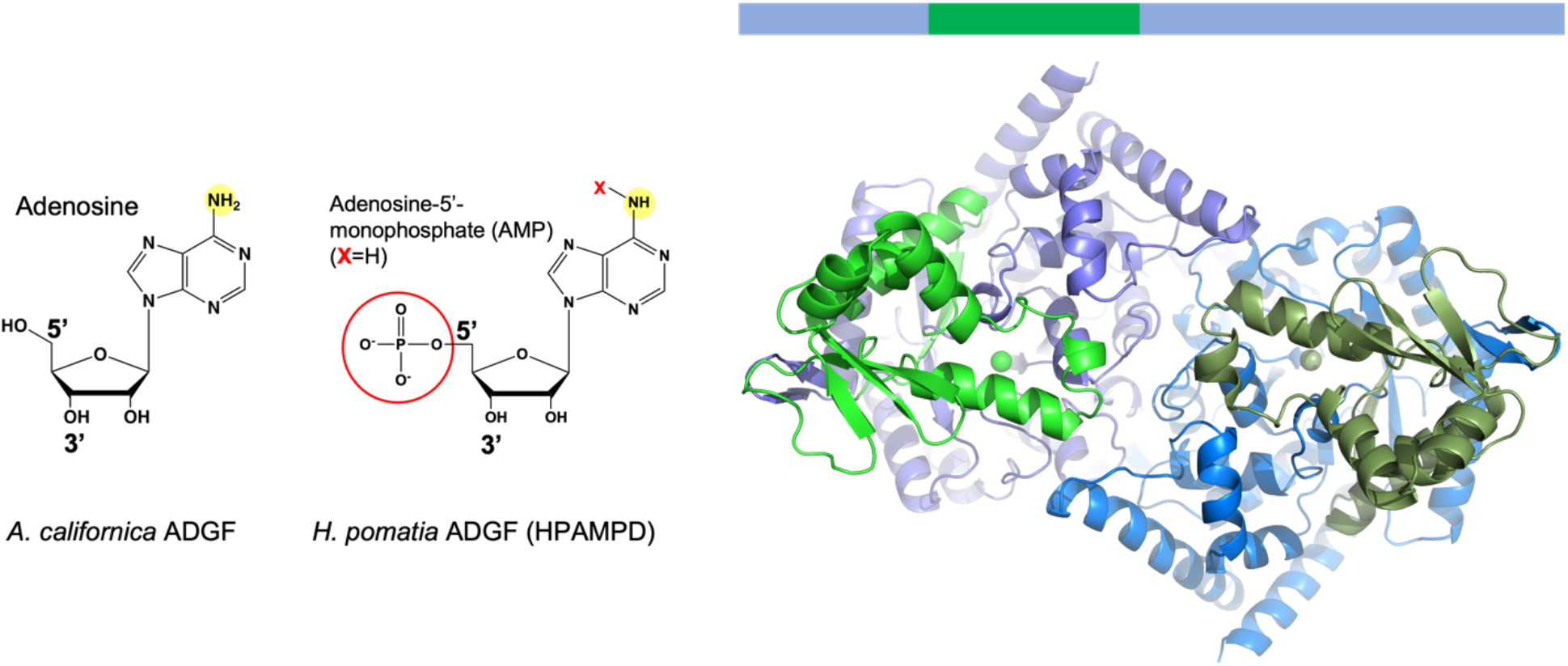

## INTRODUCTION

The <u>a</u>denosine <u>d</u>eaminase-related <u>g</u>rowth <u>f</u>actors (ADGF) family of proteins are enzymes which are understood to control the level of extracellular adenosine, an important signaling molecule that effects cellular responses ^1^. Human ADA2 is a well-studied ADGF. The concept of ADA2 as a growth factor originates from a historical study in which immature flesh fly cells exhibited an activity that stimulated insect cells growth in fresh cultured media ^2^. This secreted insect-<u>d</u>erived <u>g</u>rowth <u>f</u>actor (IDGF) was found to possess deaminase activity, which was essential for its ability to promote cell growth ^3^. Based on its protein sequence similarity to IDGF, human ADA2 was proposed to function as a growth factor ^4,5^. In this study, we focus on the deaminase activity of ADGF family members, including one identified in the sea slug, *Aplysia californica* ^6^. Based on protein sequence conservation, we recently identified a homolog of ADGF in another mollusk, *Helix pomatia* ^7^, commonly known as the Roman snail, Burgundy snail, or escargot. The *H. pomatia* deaminase shares 57% sequence identity with that of *Aplysia californica* ADGF and 38% identity to human ADA2 (Figure 1). *H. pomatia* deaminase was an unexpected member of the ADGF family, as it exhibits deaminase activity with a preference for the <u>nucleotide</u> adenosine 5’- monophosphate (AMP) over the <u>nucleoside</u> adenosine. Consequently, it was named *<u>H</u>. pomatia* <u>AMP</u> <u>d</u>eaminase (HPAMPD). Before HPAMPD, no ADGFs were known to deaminate AMP.

**Figure 1.**
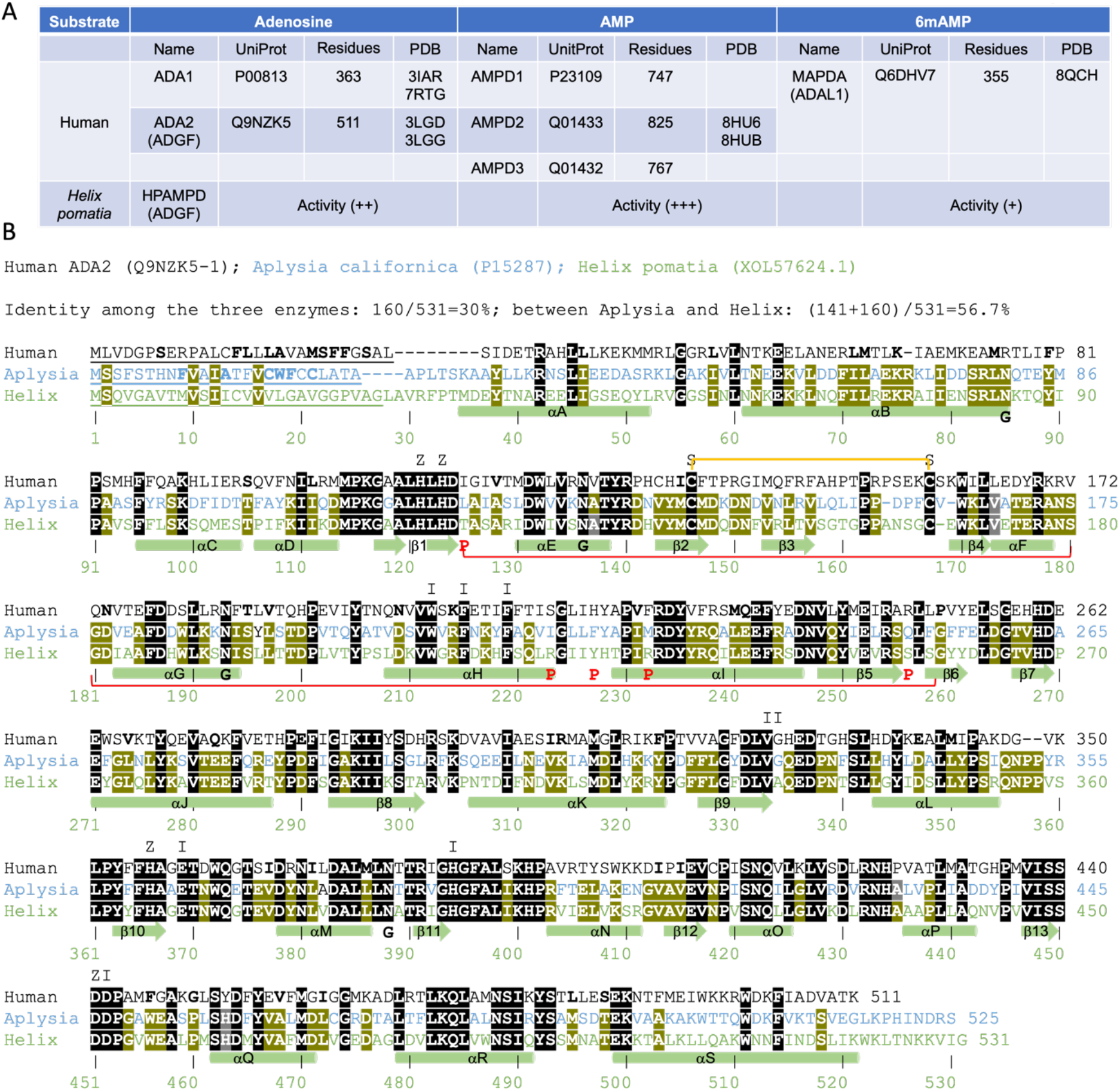
**(A)** Summary of human enzymes acting on adenosine, AMP, and 6mAMP and the relative activity with HPAMPD on each substrate. **(B)** Sequence alignment of three ADGF deaminases: human ADA2 (UniProt: Q9NZK5-1), *Aplysia* ADGF (UniProt: P15287), and HPAMPD (GenBank: XOL57624.1). The underlined N-terminal sequences represent signal peptides that control protein secretion and translocation ^65^. Invariant residues among all three enzymes are shown as white letters on a black background, while residues conserved between *Aplysia* and *Helix* ADGFs are displayed as white letters on a light green background. Secondary structure elements (α-helices and β-strands) of HPAMPD are depicted below the sequence. Letters above the human ADA2 sequence indicate conserved residues involved in zinc binding (Z) and inhibitor pentostatin binding (I). Letters P below the HPAMPD sequence marks residues uniquely involved in phosphate group binding of pentostatin 5’-monophosphate. Regions swapped between *Aplysia* ADGF and HPAMPD to generate a chimeric ADGF are indicated by a red line. A disulfide bond (S-S) between C146 and C168 is indicated by a bracket. Glycosylation at asparagine residues N85, N136, N193, and N388 are marked with the letter G (see Figure S4).

Human adenosine deaminase 1 (ADA1) deficiency, first described in the 1970s, was recognized as a cause of <u>s</u>evere <u>c</u>ombined immuno<u>d</u>eficiency (SCID) ^8^. Residual adenosine deaminase activity, attributed to adenosine deaminase 2 (ADA2), was initially studied in spleen extracts from patients with SCID and ADA1 deficiency ^9^. The ADA2 enzyme is encoded by a gene on chromosome 22, within the region affected in patients with cat eye syndrome, which led to its original gene name, <u>c</u>at <u>e</u>ye syndrome <u>c</u>ritical <u>r</u>egion 1 (*CECR1*)^10^. In contrast, ADA2 deficiency, caused by loss-of-function mutations (>100 variants) in the *ADA2*/*CECR1* gene – was only described 11 years ago. This deficiency has been linked to a broad spectrum of vascular and inflammatory phenotypes, including early-onset recurrent stroke, systemic vasculopathy, and vasculitis ^11,12^. The complexity of the phenotype and pathophysiology of ADA2 deficiency is well-documented (reviewed in ^13–18^), with an estimated carrier frequency of over 1 in 236 individuals ^19^ and more than 100 disease-associated mutations were reported ^20^. [There are 159 mutations of ADA2 in the registry of hereditary auto-inflammatory disorders mutations (Infevers) and 608 mutations of ADA2 registered in Catalogue Of Somatic Mutations In Cancer (COSMIC).]

Enzymatically, ADA1 and ADA2 catalyze the deamination of adenosine (Ado) and 2′- deoxyadenosine (dAdo), converting them to inosine and deoxyinosine, respectively. In ADA1 deficient SCID patients, plasma levels of Ado and dAdo are elevated, leading to (1) marked accumulation of dATP, derived from dAdo metabolism ^21^, and (2) increased levels of *S*-adenosyl-L-homocysteine (SAH), due to dAdo-mediated inhibition of SAH hydrolase ^22^. The excess of dATP inhibits DNA synthesis, while SAH accumulation inhibits most (if not all) transmethylation reactions catalyzed by *S*-adenosyl-L-methionine (SAM)-dependent methyltransferases, affecting multiple cellular processes. These disruptions highlight the critical role of purine metabolism and associated deaminases in cellular functions of purinergic signaling and in anti-cancer therapy resistance ^23–25^.

Human ADA1 enzyme consists of 363 residues (UniProt P00813) and adopts a single globular domain with an α/β barrel fold ^26,27^, as seen in crystal structures (human ADA1: PDB 3IAR and 7RTG; murine Ada1: PDB 2ADA) (Figure 1A). In contrast, human ADA2 (UniProt Q9NZK5; 511 residues) forms a homodimer ^5^, and features an additional ∼80 N-terminal residues responsible for dimerization as well as an extracellular signal peptide, alongside its ADA1-like catalytic domain ^28^ (PDB 3LGD and 3LGG). The presence of N-linked glycosylation, disulfide bonds, and a signal peptide at the very N-terminus indicates that ADA2 functions in the extracellular environment. In addition, ADA2 is a lysosomal protein ^29,30^.

Humans possess distinct enzymes for the deamination activity of <u>nucleosides</u> and <u>nucleotides</u>, including adenosine, adenosine 5’-monophosphate (AMP), and its methylated form, N6-methyl-AMP (6mAMP) (Figure 1A). HPAMPD exhibits deaminase activity on all three substrates, with a preference order of AMP > adenosine > 6mAMP (^7^ and this study). In contrast, human ADA2 and *Aplysia* ADGF do not deaminate AMP and 6mAMP. To investigate the substrate selectivity of HPAMPD, we determined its structures in the apo form and in complexes with pentostatin (2′-deoxycoformycin) and <u>p</u>entostatin 5’-<u>m</u>ono<u>p</u>hosphate (PMP). Additionally, we engineered a chimeric ADGF by integrating elements from *A. californica* and *H. pomatia*, resulting in a chimeric enzyme with equivalent activity on AMP and adenosine.

## RESULTS

### Structural basis of substrate specificity of the ADGFs

The human ADA2 is an adenosine deaminase with a relatively high *K*m for adenosine of 2500 μM and *k*_cat_ of 88 s^-1^ (ref. ^5^) (Figure 2A), whereas HPAMPD has a *K*m of 25 μM and a *k*cat of 170 s⁻¹ for AMP ^7^ (Figure 2B). Kinetic analyses compared the deamination rates of adenosine and AMP revealed that human ADA2 and the *Aplysia* ADGF favor adenosine over AMP by more than 100-fold and 1000-fold, respectively, as expected for ADGFs. In contrast, HPAMPD prefers AMP over adenine by 25-fold ^7^ (Figure 2C).

**Figure 2.**
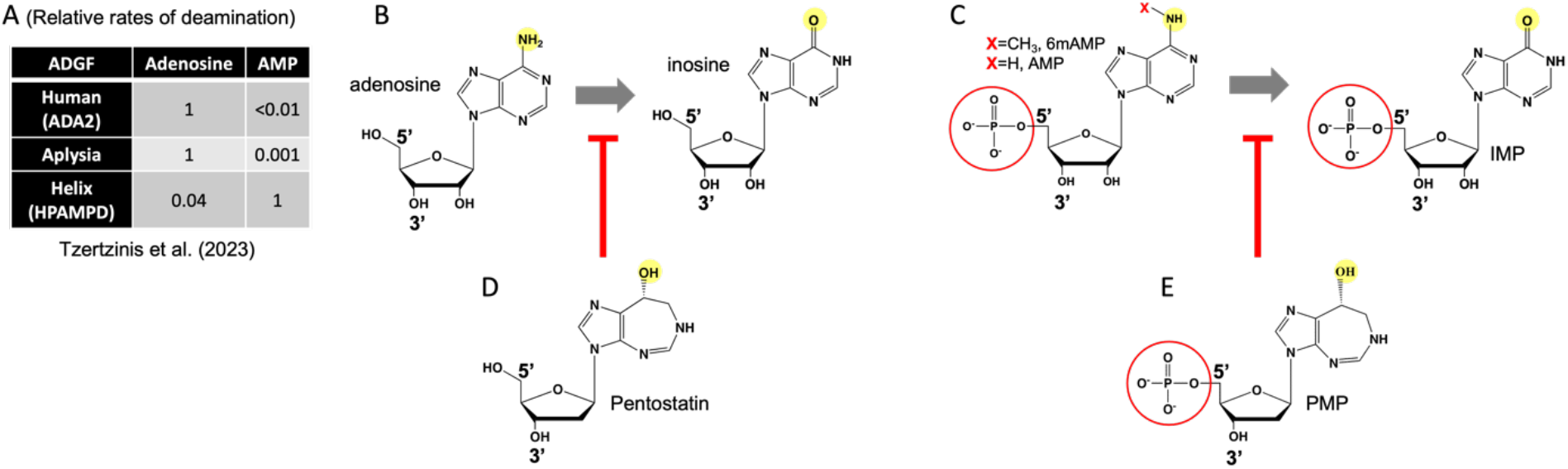
Substrate specificity of HPAMPD. (**A**) Relative rates of deamination for adenosine and AMP, adapted from ^7^. (**B**) Deamination reaction of adenosine to inosine. (**C**) Deamination reactions of AMP and 6mAMP to IMP. (**D**) Pentostatin is a transition-state analog of adenosine deamination. (**E**) PMP is a transition-state analog of AMP (and 6mAMP) deamination.

To investigate the structural basis of HPAMPD substrate selectivity, we crystallized recombinant HPAMPD, which was purified from *P. pastoris* expression ^7^. We obtained the apo-structure in the presence of zinc and also determined structures in complexes with pentostatin and pentostatin 5’-monophosphate (PMP). These structures were resolved at high resolutions ranging from 1.5-1.7 Å (Table S1). Pentostatin and its 5’-monophosphorylated form PMP are anticancer chemotherapeutic drugs that are transition state analogs of adenosine deamination ^31^ (Figure 2D-2E).

In the crystallographic asymmetric unit of space group *P*2_1_, there are two dimers of HPAMPD (molecules A and B and molecules C and D) (Figure 3A). The dimer-dimer interface is mediated mainly by water molecules (red dashed line in Figure 3A). These dimers are highly similar to each other and to the human ADA2 dimer, with a root-mean-squared deviation (rmsd) of ∼1.2 Å over 837 pairs of Cα atoms (Figure 3B). The corresponding secondary structures, consisting of 19 α-helices and 13 β-strands, are preserved throughout (Figure 1B). In the active site, the Zn ion is in a trigonal bipyramidal geometry ^32^ coordinated by three histidine residues (H121, H123, and H366) and one aspartate (D451) (Figure 3C). Additionally, a water molecule serves as the fifth ligand and acts as the attacking water nucleophile during the catalysis (Figure 3D).

**Figure 3.**
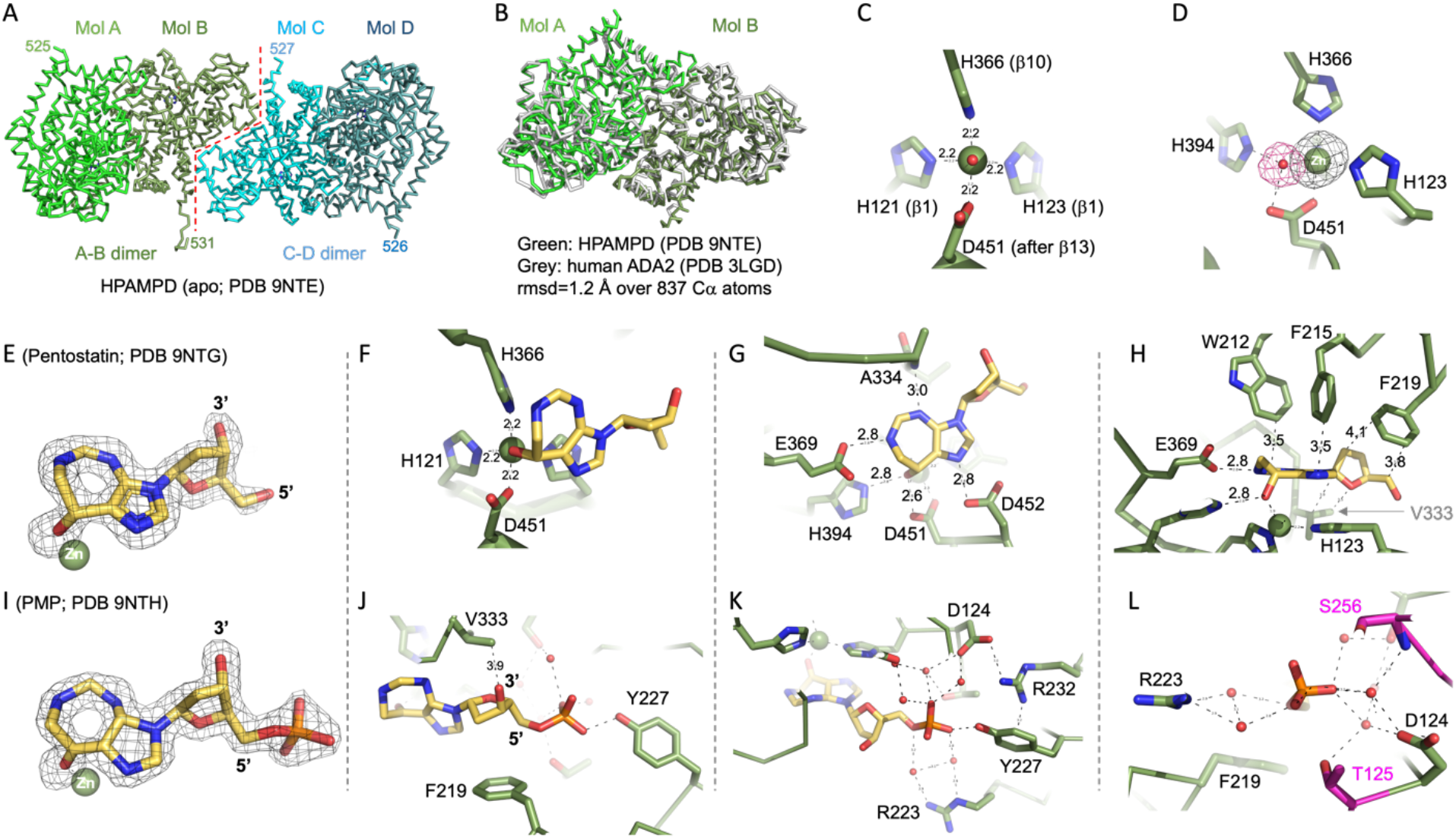
Structures of HPAMPD. (**A**) Two HPAMPD dimers in the *P*2_1_space group. (**B**) Superimposition of HPAMPD and human ADA2 dimers. (**C**) Active-site Zn coordination viewed from the Zn-bound water molecule (colored in red). (**D**) Omit electron densities contoured at 5α for Zn and 5α for the water. (**I**) Omit electron density contoured at 5α for bound pentostatin. (**E**) The hydroxyl group of the coformycin ring occupies the position of the attacking water molecule. (**F**) Hydrogen bonds formed by the coformycin ring. (**G**) Aromatic and hydrophobic interactions stabilizing pentostatin binding. (**H**) Omit electron density contoured at 5α for bound PMP. (**I**) Direct hydrogen bond between the phosphate group of PMP and Y227. (**J**) R232 forms a bridge between D124 and Y227. (**K**) Water-mediated interactions involving the phosphate group of PMP.

In the complex structure with pentostatin (Figure 3E), the hydroxyl group of the seven-membered coformycin ring occupies the position of the attacking water (Figure 3F). The configuration mimics the transition state generated by the zinc-activated hydroxyl attack on the face of adenine ring. The nitrogen atoms of the coformycin ring establish hydrogen bonds with the carboxylate groups of E369 and D452, as well as the main-chain amide group of A334 (Figure 3G). Additionally, the coformycin ring is stabilized through aromatic interactions, being sandwiched between W212 and F215 from above and H123 from below (Figure 3H). The ribose ring is positioned between F219 and V333. All residues involved in pentostatin binding are either identical or highly conserved among the three enzymes: human ADA2, *Aplysia* ADGF, and HPAMPD (Figure 1B).

Next, we examined the structure of HPAMPD bound to pentostatin 5’-monophosphate (PMP) (Figure 3I). All interactions observed with pentostatin are retained in PMP. The additional interaction involving the 5’-monophosphate group includes a direct hydrogen bond with the hydroxyl group of Y227 (Figure 3J), which is stabilized by R232 through a bridge with D124 (Figure 3K). Additionally, a network of water-mediated hydrogen bonds is formed with the side chains of D124, T125, R223 and S256 (Figure 3L). Among them, except for the invariant D124, five polar and positively charged residues are unique to HPAMPD involved in phosphate group binding (via direct H-bond or indirect water-mediated interactions) – T125 after β1, R223 from αH, Y227 located in the loop between αH and αI, R232 from αI and S256 after β5 (Figure 1B). In contrast, the corresponding residues are hydrophobic in *Aplysia* ADGF (Leu, Ile, Phe, Met and Phe), and in human ADA2 they are Ile, Ser, His, Phe, and Leu (Figure 1B). These hydrophobic and/or smaller residues are less suited for electrostatic interactions with the negatively charged phosphate group, highlighting a structural adaptation unique to HPAMPD.

The 3’ hydroxyl (OH) group of the ribose moiety is in van der Waals contact with V333 (Figure 3J). The addition of a 3’-monophosphate creates a repulsive force with V333 (Figure S1A), leading to the loss of HPAMPD activity on 3’-AMP, 3’,5’-adenosine diphosphate, and 3’,5’-cyclic AMP ^7^. In contrast, the extension of 5’-AMP (100% activity) to 5’-ADP (74-95% activity) preserves function ^7^ by orienting the additional 5’-phosphate group away from the active site and toward the solvent (Figure S1B). Additionally, HPAMPD maintains moderate activity (25-36%) on 5’-ATP ^7^ (Figure S1C).

### Generation of chimeric ADGF with altered activity

To clarify the origin of substrate selectivity, we hypothesized that residues near the phosphate group were responsible for binding PMP. To test this, we first generated Chimera-1 by replacing two short segments within the backbone of *Aplysia* ADGF with the corresponding HPAMPD residues: the six-residue sequence immediately after β1: L<u>A</u>I<u>A</u>SL in *Aplysia* was substituted with T<u>A</u>S<u>A</u>RI (residues 125-130 of HPAMPD), and the eight-residue sequence of strand β5 and its adjacent loop: I<u>E</u>L<u>RS</u>Q<u>L</u>F in *Aplysia* was replaced with V<u>E</u>V<u>RS</u>S<u>L</u>S (residues 251-258 of HPAMPD). These sequences differ between the two enzymes, with 4 out of 6 and 4 out of 8 residues diverging in each segment, respectively, including residues T125 and S256, which are involved in water-mediated phosphate interactions (Figure 3K). However, the chimeric *Aplysia* enzyme (Chimera-1) did not recognize AMP as a substrate. With adenosine as substrate, it exhibited activity similar to that of wild type *Aplysia* ADGF, which was more than 100 times higher than its activity on AMP (Figure 4A).

**Figure 4.**
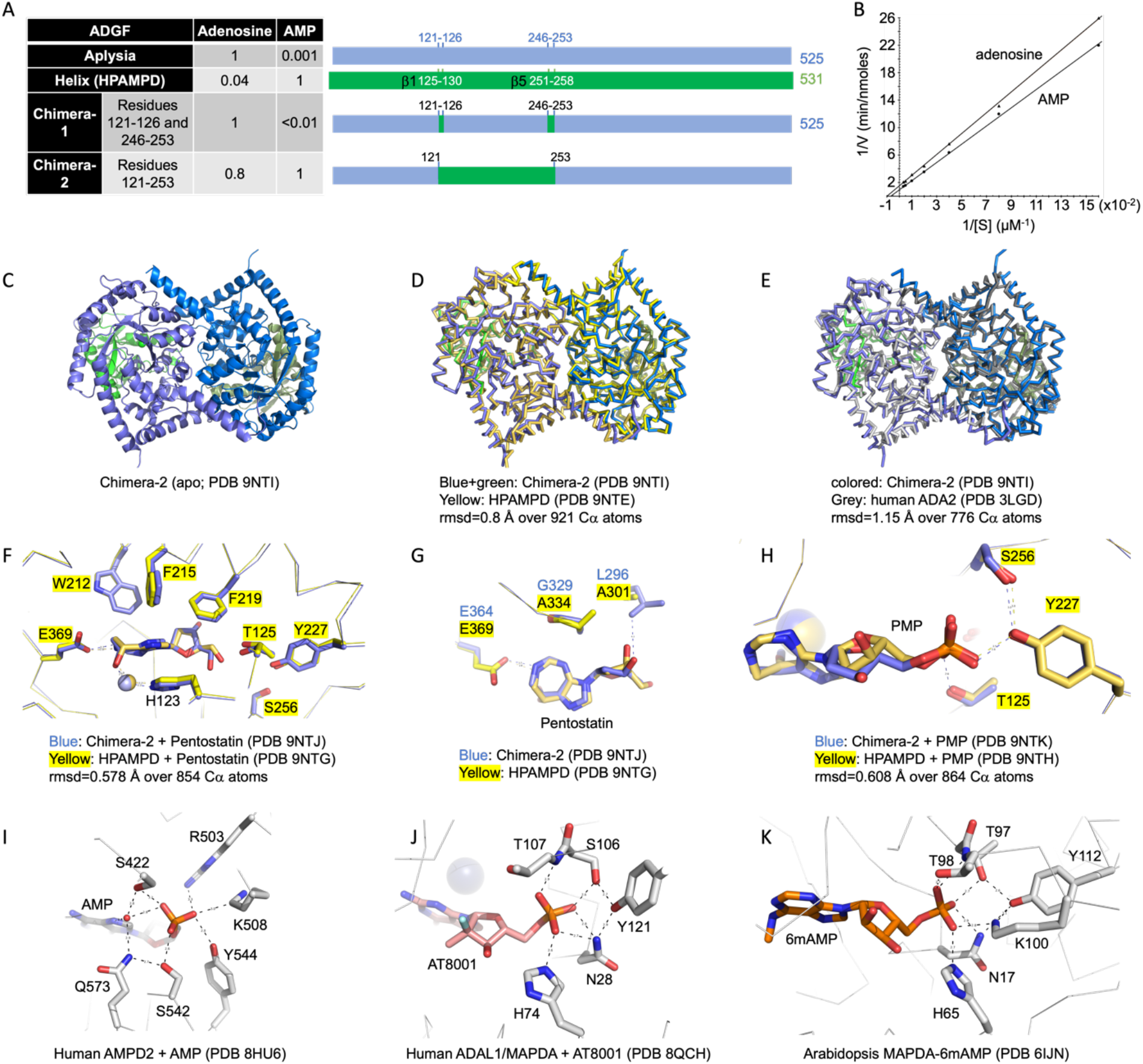
Substrate specificity of engineered chimera ADGF. (**A**) Relative deamination rates of for 100 µM adenosine and AMP for engineered chimera enzymes, along with wild-type *Aplysia* and *Helix* ADGF, depicted in cartoon representations to the right. (**B**) Enzymatic activities of the Chimera-2 variant. (**C**) Dimeric structure of Chimera-2. (**D**) Superimposition of Chimera-2 and HPAMPD dimers. (**E**) Superimposition of Chimera-2 and human ADA2 dimers. (**F**) Bound conformation of pentostatin in the active sites of Chimera-2 and HPAMPD. (**G**) Interactions between pentostatin and *Aplysia* ADGF-specific residues (L296 and G329 colored in blue). (**H**) Bound PMP in the active sites of Chimera-2 and HPAMPD. (**I**) Phosphate interaction within the AMP-bound active site of human AMPD2 (PDB 8HU6). (**J**) Phosphate interaction within the active site of human ADAL1/MAPDA bound with an AMP analog (PDB 8QCH). (**K**) Phosphate interaction within the active site of *Arabidopsis* MAPDA with a bound 6mAMP (PDB 6IJN)

Next, we replaced the entire region (from L<u>A</u>I<u>A</u>SL through I<u>E</u>L<u>RS</u>Q<u>L</u>F) with residues 125-258 of HPAMPD, creating Chimera-2. The engineered Chimera-2 efficiently deaminates AMP with a *K*_M_ of 135 µM and *k*_cat_ of 33 s^-1^ and deaminates adenosine with a *K*_M_ of 104 µM and *k*_cat_ of 22 s^-1^ (Figure 4B), demonstrating altered substrate specificity. Notably, Chimera-2 retains the characteristic of adenosine deamination activity of *Aplysia* ADGF. To explore potential structural differences, we crystallized Chimera-2 in its apo form as well as in complexes with pentostatin and PMP (Table S1).

Like human ADA2, Chimera-2 adopts a dimeric structure (Figure 4C). As expected, Chimera-2 closely resembles HPAMPD (rmsd=0.44 Å) and human ADA2 (rmsd=1.15 Å) in overall structures (Figure 4D-4E). The bound pentostatin has a similar conformation in the active sites of both Chimera-2 and HPAMPD, with all conserved residues aligning perfectly (Figure 4F). Additionally, some interactions occur with *Aplysia* ADGF-specific residues, such as Leu296 (which corresponds to Ala301 in HPAMPD) (Figure 4G). The binding mode of PMP is also identical between Chimera-2 and HPAMPD, including the coordination of the phosphate group by polar residues (Figure 4H).

In addition to ADA1 and ADA2, humans have three AMP deaminase genes: AMPD1 expressed in muscle (UnitProt: P23109), AMPD2 in liver (Unit Prot: Q01433), and AMPD3 in erythrocytes (UnitProt: Q01432) ^33^ (Figure 1A). The AMPD enzymes are significantly larger in size (comparing 825 residues in AMPD2 and 511 residues in ADA2), yet they share a conserved active site. In a recent structure of AMPD2 bound to AMP ^34^, the negatively phosphate group of AMP is surrounded by positively charged residues (R503 and K508), polar residues (S422, S542 and Y544), and ordered water molecules (Figure 4I).

Moreover, X-ray structures are available for 6mAMP deaminases, including human ADAL1 ^35^ and *Arabidopsis* homolog of MAPDA ^36,37^. In both cases, a set of polar residues (Ser, Thr, Asn, His and Tyr), along with a positively charged lysine, interact with the monophosphate group of either the guanosine analog AT8001 (Figure 4J) or the substrate 6mAMP (Figure 4K). Although these phosphate-interacting residues are not perfectly conserved in primary sequences (Figure S2 and S3), the four enzymes compared - HPAMPD, human AMPD2, human ADAL1, and *Arabidopsis* MAPDA - utilize similar interactions for AMP binding. These residues are positioned in conserved structural regions, including the loop immediately following strand β1, the loop between helices αH and αI, and the end of strand β5 and its adjacent loop (Figure S2 and S3). This structural similarity is evident only through structured-based alignments due to large insertions and deletions between secondary structural elements β1 to β5. As demonstrated by our Chimera-2 enzyme, the entire β1 to β5 region of HPAMPD, containing all three structural elements, is essential for AMP binding and activity.

### Discrimination between AMP and 6mAMP

While the deamination of AMP to generates IMP and ammonia, the deamination of N6-methyl- AMP (6mAMP) also generates IMP and methylamine (Figure 3B). In human, a specific 6mAMP deaminase (MAPDA), also known as ADA-like protein 1 (ADAL1) (UnitProt: Q6DHV7), carries out this reaction (Figure 1A) ^38^. The *Arabidopsis* homolog of MAPDA is highly specific for 6mAMP and does not act on adenosine, N6-methyladenosine, AMP, N6-methyl-ATP and O6-methyl-guanosine ^39^. Interestingly, we found that HPAMPD is also capable of deaminating 6mAMP, albeit at a much slower rate. The relative activity for AMP, adenosine, and 6mAMP follows a ratio of 1: 0.04: 0.013 (Figure 5A). In Chimera-2 the residues surrounding the N6 position of adenine ring have not been altered. Despite this, the 10-fold difference in 6mAMP activity between Chimera-2 and HPAMPD is not fully explained by the current structural data. The deamination rate of 6mAMP in Chimera-2 is 1000-fold slower than that of AMP (Figure 5A).

**Figure 5.**
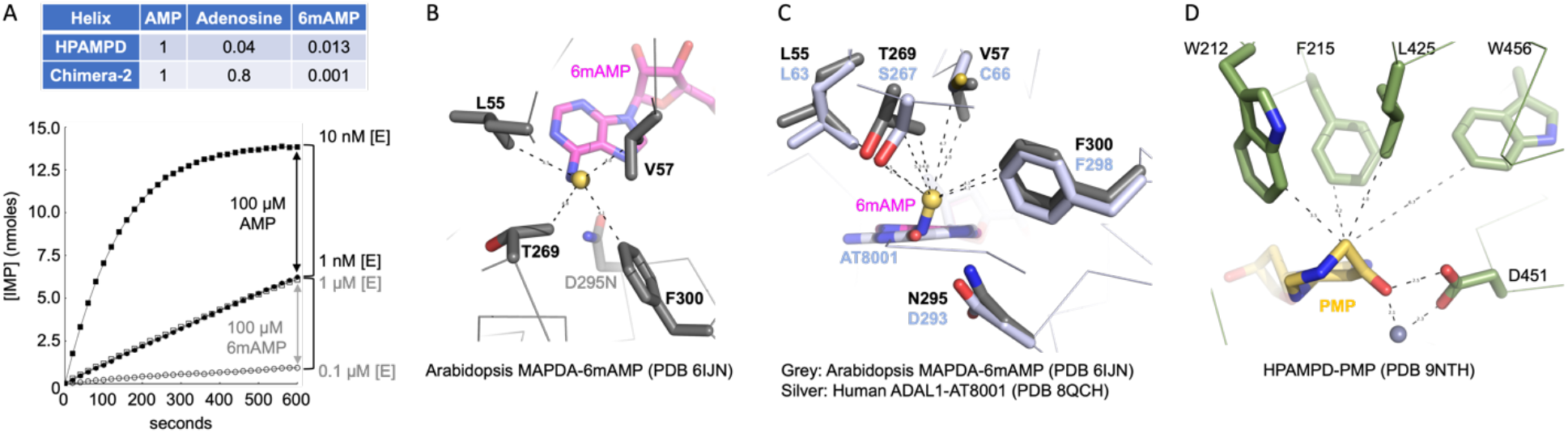
HPAMPD and Chimera-2 have weak activity on 6mAMP. (**A**) Relative deamination rates of AMP, adenosine, and 6mAMP by HPAMPD and Chimera-2. The graph displays the different deamination rates of Chimera-2 on AMP and 6mAMP at two enzyme concentrations. (**B**) Binding of the N6-methyl group (in yellow sphere) of 6mAMP in the inactive mutant (D295N) of *Arabidopsis* MAPDA (PDB 6IJN). (**C**) Superimposition of human ADAL1 (PDB 8QCH) and its *Arabidopsis* homolog (PDB 6IJN). (**D**) HPAMPD, which exhibits low activity on 6mAMP, and human AMPD2, which has no known activity on 6mAMP, share three large aromatic residues and one leucine that form an aromatic cage.

In the structure of the catalytically inactive mutant (D295N) of the *Arabidopsis* MAPDA homolog ^36^, which permits substrate 6mAMP binding without undergoing catalysis, the N6-methyl group is positioned out of plane with the adenine ring, oriented opposite to the attacking nucleophile, and forms van der Waals contacts with residues L55, V57, T269, and F300 (Figure 5B). The corresponding residues in human ADAL1 are L63, C66, S267, and F298 (Figure 5C). Although only two of these four residues (Leu and Phe) are conserved, the variation (Val vs. Cys and Thr vs. Ser) involve residues of similar size. In contrast, three of four corresponding residues in HPAMPD - W212, F215, L425, and W456 - are larger aromatic residues, which may contribute to its low activity on 6mAMP (Figure 5A). These residues form an aromatic pocket surrounding the C9 carbon atom of the non-planar coformycin ring (Figure 5D). Similarly, in human AMPD2, which has no known activity on 6mAMP, the equivalent residues consist of three phenylalanines (F433, F436, F715) and one leucine (L686) (Figure S2). We propose that the size of these four residues may play a key role in determining deaminase activity on 6mAMP.

## DISCUSSION

Adenosine deaminase deficiency is a known cause of <u>s</u>evere <u>c</u>ombined immuno<u>d</u>eficiency (SCID)^40,41^. Humans possess at least six adenylate deaminases, - ADA1/2, AMPD1-3, and MAPDA/ADAL1 – each with distinct substrate specificities for adenosine, AMP, and 6mAMP, respectively (summarized in Figure 1A). While no known methyltransferase modifies AMP, N6-methylation of adenine in specific RNA sequences is the most common internal mRNA modification in eukaryotes ^42,43^. When 6mA-containing RNA is degraded in the cytoplasm, 6mAMP is released. The deamination of 6mAMP may prevent its conversion to 6mADP and eventually 6mATP, which could otherwise be randomly incorporated into newly synthesized RNA molecules by RNA polymerase ^39,44^.

Interestingly, the foot muscle of the mollusk *H. pomatia* is one of the richest sources of “non-specific” adenylate deaminase(s), exhibiting activities on AMP, ADP, ATP and NADH ^45,46^. The recombinant form of HPAMPD shares a similar substrate spectrum to the native enzyme, which has been purified at least twice from *H. pomatia* foot muscle - first in 1983 ^45^ and again in 2023 ^7^. Structurally, HPAMPD adopts a fold similar to human ADA2, a nucleoside adenosine deaminase, but has acquired the ability to interact with the 5’-monophosphate group through polar and charged residues located in three key structural elements: (1) the loop immediately following strand β1, (2) the loop between helices αH and αI, and (3) the end of strand β5 and its adjacent loop. The phosphate-binding feature is characteristic of nucleotide deaminases, such as human AMP and 6mAMP deaminases. We note that the *Aplysia* ADGF discriminates by 2000-fold against adenosine when the N6 position is methylated (6mAMP), while HPAMPD discriminates only about 100-fold against 6mAMP. Interestingly, Chimera-2 while accepting the phosphorylated substrate, which is a property of HPAMPD, discriminated by a 1000-fold against 6mAMP, a property similar to that of the *Aplysia* ADGF towards deamination of 6mAMP.

As mentioned above, the deamination of AMP (or its 2’-deoxy form, dAMP) and 6mAMP (or its 2’-deoxy form, 6m(d)AMP) produces the common inosine derivatives IMP (or dIMP). A recent study suggested that dIMP may serve as a precursor for immune messengers in bacterial anti-phage defense ^47^. In *Escherichia coli* NCTC13216, dIMP production begins with the deamination of dAMP by a host bacterial enzyme called KomA, which works in conjunction with deoxynucleotide phosphate kinases to generate dIDP and ultimately dITP as signaling molecules^47^. KomA, a small protein of 326 amino acids, shares the highest sequence similarity with human ADAL1 (Figure S5A). The AlphaFold-predicted structure ^48^ of KomA aligns closely with experimentally determined structures of human ADAL1 and *Arabidopsis* MAPDA, with an RMSD of 1.7-1.8 Å (Figure S5B-D). While KomA exhibits *in vitro* activity on both 2’-hydroxy and 2’- deoxy forms of AMP and ADP ^47^, its high structural similarity and comparable size to 6mAMP deaminases (ADAL1/MAPDA) suggest that its activity on 6mAMP, 6m(d)AMP as well as potential activity on DNA or RNA, still needs to be investigated. Notably, both bacterial and phage DNA can undergo extensive adenine hypermethylation, primarily as part of the restriction-modification defense system that protects against exogenous DNA ^49–52^. This widespread adenine methylation may have functional implications for KomA-mediated immune defense. Recently the deamination of deoxyadenosine in DNA in vitro was described for human ADA2, potentially involved in regulating immune sensing of DNA ^30^.

## Material and methods

### Protein purification

The HPAMPD and *Aplysia* ADGF proteins were prepared as described ^7^. The Chimera-1 and Chimera-2 ADGFs used in this study were all expressed in *Pichia pastoris* by integrating an expression cassette coding for each and were cultured as described ^7^. The Chimera-1 protein was directly assayed from the clarified spent culture medium.

*P. pastoris* harboring the integrated Chimera-2 expression cassette was cultured in 2400 ml of Tolner media ^53^ with 1% glycerol and 100 μg/ml ampicillin at 30° C. The secreted expression of AMP deaminase activity was measured over time of culturing until culture fluid expressed highest level of secreted AMP deaminase activity. At the time of the highest expression the culture was clarified by centrifugation and the supernatant culture fluid was diluted 2-fold with 20 mM Na-acetate pH 6.0, applied to a Heparin Hyper D column and eluted with a gradient from 50 mM to 1 M NaCl. The peak of AMP deaminase activity was diluted 8-fold into 20 mM Tris-HCl and applied to a HiTrap^TM^ Q column. Activity was eluted with a gradient from 50 mM to 1 M NaCl. The peak of activity was pooled and was treated with a recombinant protein fusion of Endoglycosidase H and maltose binding protein (Endo H_f_, NEB) by first dialyzing against 20 mM Na acetate pH 6.0, 1 mM EDTA, 50 mM NaCl, 1 mM DTT. Endo H_f_ was added to the dialysate and incubated at 25° C overnight. Endo H_f_ was removed from the preparation by passing the supernatant through amylose resin (NEB). The flow-through was applied to a HiTrap^TM^ SP column. The flow-through from the SP column was diluted threefold in 10 mM Tris-HCl (pH 7.5) before being loaded onto a HiTrap^TM^ Q column. The peak of Chimera-2 activity was pooled and dialyzed against 50% glycerol, 10 mM Tris-HCl pH 7.4, 1 mM DTT, 0.1 mM EDTA. The Chimera-2 preparation is stored at -20° C prior to use.

### Construction of Chimera-1 and Chimera-2

The plasmid carrying the HPAMPD is described ^7^. The plasmid PD912-GAP carrying the *Aplysia* ADGF was constructed by Genscript (Piscataway, NJ) (Table S2 for plasmid sequence). The chimeras were constructed from two fragments that were generated by PCR. One linear fragment was common to both constructs and was generated by PCR using the *Aplysia* ADGF plasmid as template with primers TRA and EFA. The other fragment which was specific for Chimera-1 was generated by PCR using the same template with primers TFA and ERA (Table S2). The two fragments for Chimera-1 were assembled with NEBuilder HiFi DNA Assembly. The fragment specific for Chimera-2 was generated by PCR using the HPAMPD plasmid as template with primers TFH and ERH. Chimera-2 was assembled from the common fragment and Chimera-2 specific fragment with NEBuilder HiFi DNA Assembly. Construction of the expression cassettes and the transformation of *P. pastoris* followed the strategy described ^7^.

### Deaminase Activity and binding Assays

The assay is based on the classic Kalckar spectrophotometric method ^54^, which measures the decrease in optical density at 265 nM. The reaction buffer is 20 mM Tris-HCl at pH 7.5, 50 mM NaCl and the substrate concentration is 100 μM (AMP or adenosine or 6mAMP). The reaction volume ranged from 200 to 400 μL. The optical density measurement was determined using a SpectraMax M3 spectrophotometer.

### Crystallography

Purified HPAMPD, originally in 50% glycerol, 10 mM Tris pH 7.4, 1 mM DTT and 0.1 mM EDTA was buffer exchanged to 10 mM Tris pH 7.4, 50 mM NaCl and concentrated to ∼77 µM. HPAMPD (50 µM) was incubated with and without pentostatin or PMP in 1:3 molar ratio (up to 150 µM pentostatin or PMP) independently in buffer 10 mM Tris pH 7.4, 50 mM NaCl for 30 min at room temperature and crystallization trays were setup using sitting drop vapor diffusion method using commercial screens, Crystals of apo HPAMPD and in complex with pentostatin were observed in 0.1 M BIS-TRIS pH 6.5 and 20% (w/v) polyethylene glycol (PEG) monomethyl ether 5000 whereas crystals of HPAMPD with PMP were observed in 0.1 M HEPES pH 7.5 and 25% (w/v) PEG 3350 within 1 day of setup. All crystals were soaked in cryoprotectant containing 20% ethylene glycol supplemented in the mother liquors followed by flash freezing in liquid nitrogen.

Purified Chimera-2 was diluted 5 times in buffer [20 mM Tris pH 7.4, 100 mM NaCl, 1 mM DTT, 0.1 mM EDTA] and concentrated to ∼10 mg/ml (∼85 μM). Crystallization screens of purified Chimera-2 protein were setup at 2.5, 5 and 10 mg/ml using commercial crystallization screens (Hampton research). Needle shaped crystals of apo Chimera-2 were observed at all three concentrations tested in multiple conditions within 24 h, and crystals obtained in 0.1 M sodium acetate trihydrate pH 4.5, 25% PEG3350 resulted in the best dataset.

Chimera-2 enzymes (50 µM) were mixed with pentostatin or PMP (150 µM each) in 1:3 molar ratio independently in buffer 20 mM Tris pH 7.4, 100 mM NaCl, 1 mM DTT for 30 min at room temperature and crystallization screens were setup using sitting drop vapor diffusion method (0.2 µl protein solution +0.2 µl reservoir solution). Needle shaped crystals of Chimera-2+pentostatin and Chimera-2+PMP were observed in multiple conditions within 24 h. The conditions were further optimized by varying the percentage of PEG 1500 (15-25%) and glycerol (5-25%) and crystallization drops were setup manually. The best crystals of Chimera-2+pentostatin were obtained in 15% glycerol and 25% PEG 1500 after 15 days of setup and for Chimera-2+PMP were obtained in 0.2 M magnesium chloride hexahydrate, 0.1 M BIS-TRIS pH 5.5 and 25% (w/v) PEG 3350 after 2 days.

Diffraction data were collected at the AMX beamline 17-1D-1 of NSLS-II at Brookhaven National Laboratory and SERCAT beamline 22ID of APS at Argonne National Laboratory. Data were processed with HKL2000 ^55^ and with DIALS ^56,57^ and AIMLESS ^58^. All the crystal structures were solved by molecular replacement using either search model of human ADA2 (PDB 3LGG) ^28^ or an AlphaFold ^48^ predicted model from COLAB server ^59^. Phenix Refine ^60^ was used for all the rounds of refinements and the quality of the structure was improved manual inspection and model (re)building with COOT ^61,62^ along with iterative rounds of Phenix refinements. All the structures were validated by PDB validation server ^63^ and images were generated using Open-Source PyMOL molecular graphics system, version 3.0.3, Schrodinger, LLC. Symmetry related molecules were identified, and their dimer interface areas were calculated using PISA webserver at the European Bioinformatics Institute ^64^.

## Data Availability

The protein sequence of HPAMPD has been deposited to GenBank with accession number XOL57624.1. The X-ray structures (coordinates and structure factor files) reported here have been deposited to PDB and are publicly available as of the date of publications. PDB accession numbers are 9NTE and 9NTF (HPAMPD in the apo form), 9NTG (HPAMPD in complex with pentostatin), 9NTH (HPAMPD in complex with PMP), 9NTI (Chimera-2 in the apo-form), 9NTJ (Chimera-2 in complexes with pentostatin), and 9NTK (Chimera-2 in complex with PMP).

## Acknowledgements

We thank Daniel Kneller, Larry McReynolds, and Zhi-Yi Sun of New England Biolabs (NEB) for comments. We thank M. Wulf (NEB) for providing PMP and L. Liang (NEB) for P. pastoris fermentations. We thank X. Kong (New York University) for assistance of access to 17-ID-1 beamtime. We thank the beamline scientists at the National Synchrotron Light Source II, Brookhaven National Laboratory and the Southeast Regional Collaborative Access Team (SER-CAT) at the Advanced Photon Source (APS), Argonne National Laboratory, USA. The use of SER-CAT is supported by its member institutions and equipment grants (S10_RR25528, S10_RR028976, and S10_OD027000) from the US National Institutes of Health. Use of the APS was supported by the U.S. Department of Energy, Office of Science, Office of Basic Energy Sciences, under contract W-31-109-Eng-38. The work was supported by New England Biolabs (to I.S.), U.S. National Institutes of Health grant R35GM134744 (to X.C.), and Cancer Prevention and Research Institute of Texas grant RR160029 (to X.C., who is a CPRIT Scholar in Cancer Research).

## Author contributions

G.T. constructed chimeras. I.S. performed protein purifications and deaminase assays. G.K. performed crystallization and substrate modeling. G.K. and J.R.H. performed X-ray crystallography experiments. J.Z. performed the phylogenetic tree analysis and AlphaFold prediction of KomA. I.S. initiated the study, and I.S. and X.C. organized and designed the scope of the study.

