## Supplementary figures and tables for "Structural basis of substrate specificity of *Helix pomatia* AMP deaminase and a chimeric ADGF adenosine deaminase"

**5** Figures and **2** tables

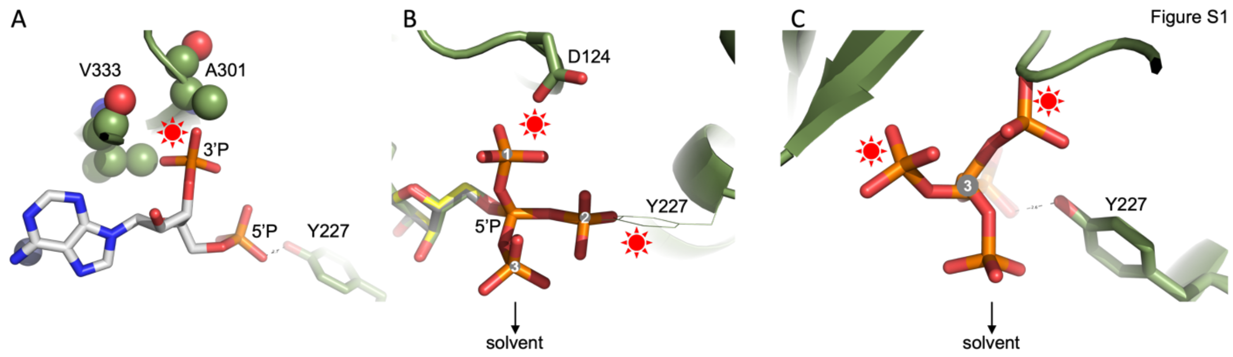


**Figure S1**. (**A**) Model of 3',5'-adenosine diphosphate, showing that the 3'-monophosphate group clashes into two hydrophobic residues of A301 and V333 in HPAMPD. (**B**) Model of 5'-ADP, illustrating that the second 5'-phosphate group can adopt three alternative conformations: conformation 1 clashes with the negatively charged D124, conformation 2 clashes with Y227, conformation 3 is positioned toward the solvent, allowing accommodation without disrupting the existing structure of HPAMPD. (**C**) Model of 5'-ATP, where the third phosphate group is added onto the conformation 3 of ADP. The outermost phosphate group is oriented toward the solvent, making it accommodable within the existing structure.

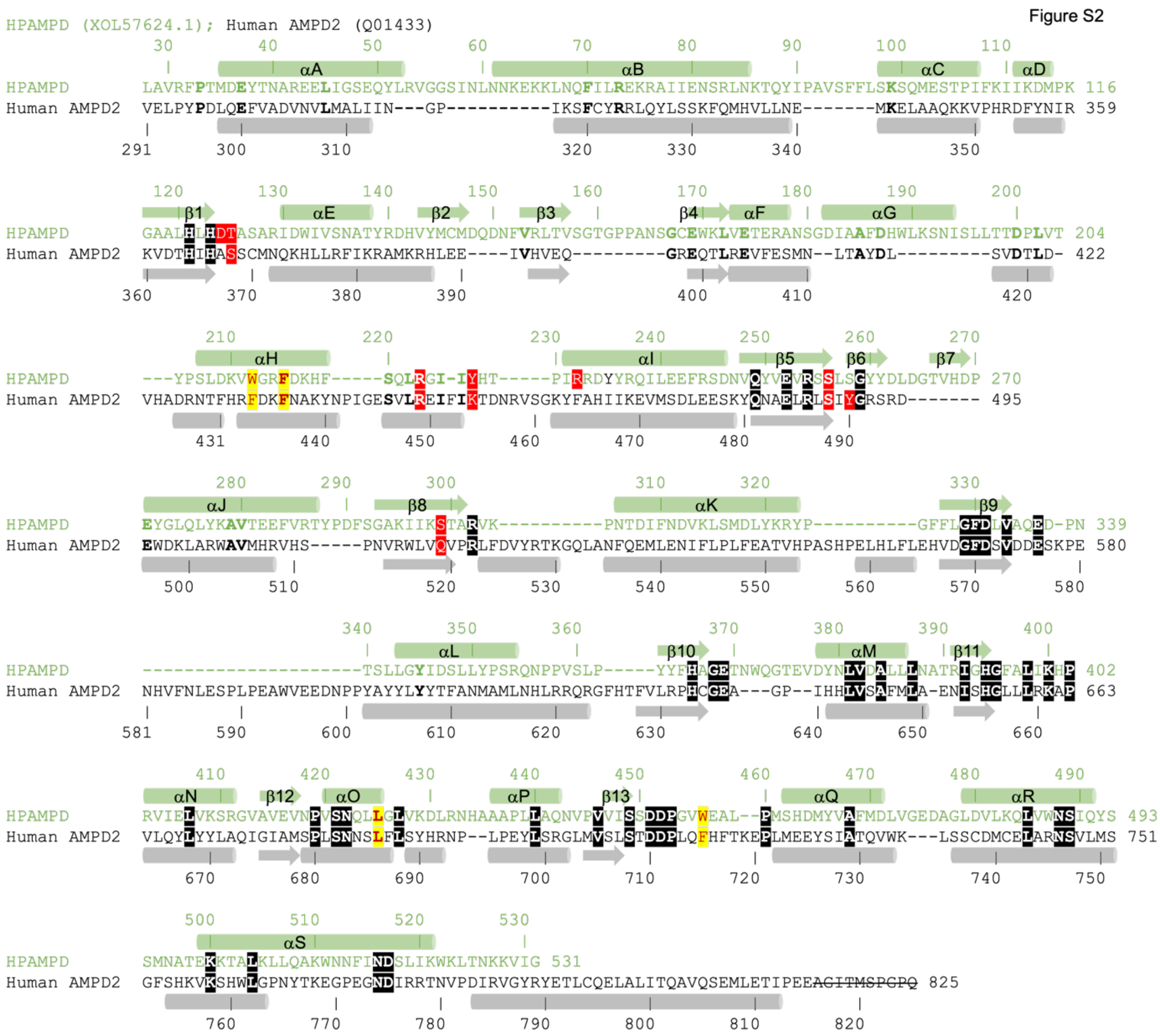


**Figure S2**. Structure-based sequence alignment of two AMP deaminases: HPAMPD (GenBank: XOL57624.1) and human AMPD2 (UniProt: Q01433). Human AMPD2 contains an additional ~290 N-terminal residues that are absent in HPAMPD. Corresponding structural elements are indicated above or below the sequences, with gaps introduced based on structural alignment. Structurally aligned conserved residues are highlighted as white letters on a black background. Residues involved in 5'-monophosphate binding are shown as white letters on a red background, while residues forming the aromatic cage (see Figure 5D) are highlighted with a yellow background. Crossed-out C-terminal residues in human AMPD2 is disordered in the structure.

Figure S3

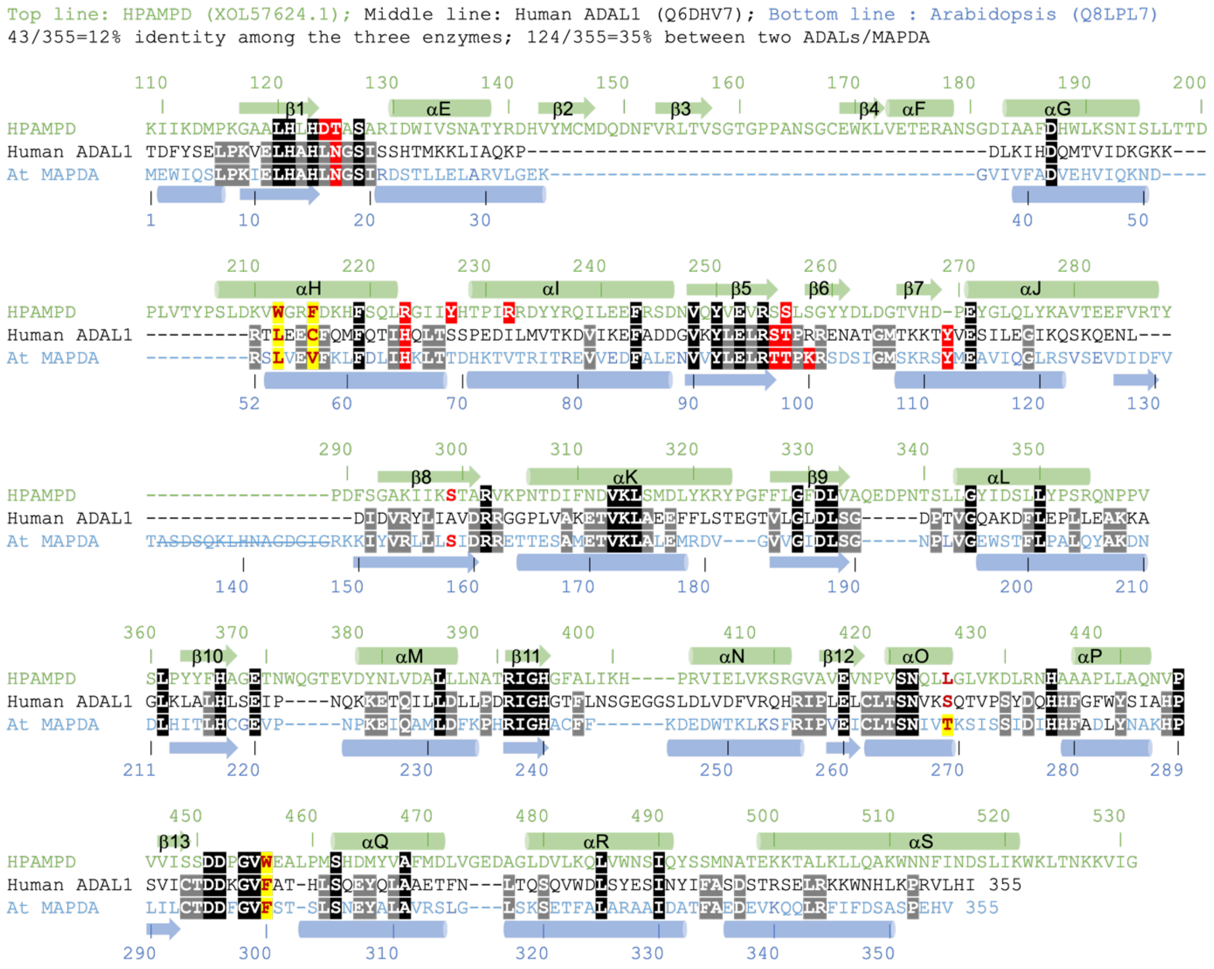


**Figure S3**. Structure-based sequence alignment of HPAMPD (GenBank: XOL57624.1) and two 6mAMP deaminases: human ADAL1 (UniProt: Q6DHJV7) and *Arabidopsis* MAPDA (UniProt: Q8LPL7). HPAMPD contains an additional ~110 N-terminal residues that are absent in human ADAL1 and *Arabidopsis* MAPDA. Corresponding structural elements are indicated above or below the sequences, with gaps introduced based on structural alignment. Structurally aligned conserved residues are highlighted as white letters on a black background. Residues involved in 5'-monophosphate binding are shown as white letters on a red background. Residues interacting with the N6-methyl group of 6mAMP in *Arabidopsis* MAPDA (see Figure 5B) and human ADAL1 (see Figure 5C), as well as the corresponding residues in HPAMPD (see Figure 5D), are highlighted with a yellow background. Crossed-out residues (133-147) in *Arabidopsos* MAPDA correspond to a disordered region in the structure and are absent in HPAMPD and human ADAL1.

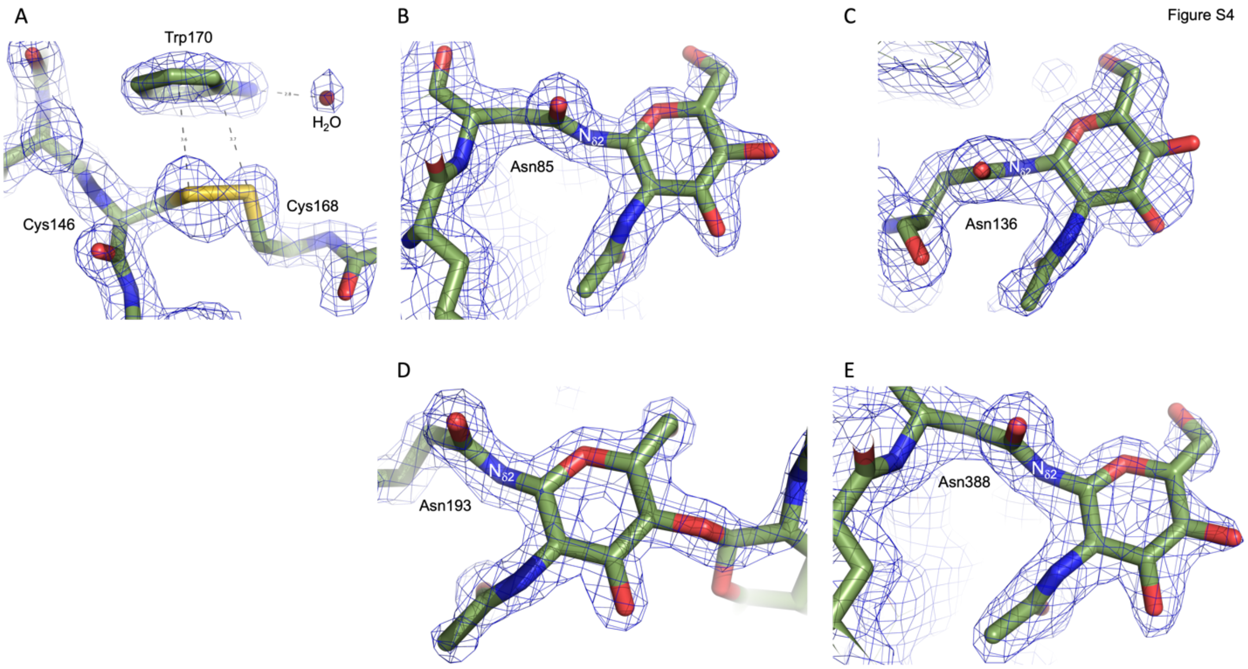


**Figure S4**. Examples of electron densities in the HPAMPD-PMP complex structure (PDB 9NTH). The 2Fo-Fc electron density is contoured at 1.5σ above the mean. (**A**) Cys146 and Cys168 forms a disulfide bond. (**B**) A single modification of *N*-acetylglucosamine (NAG) is attached to the side chain Nδ2 atom of Asn85. (**C**) A single modification is attached to the side chain Nδ2 atom of Asn136. (**D**) Two modification units are attached to the side chain Nδ2 atom of Asn193, with incomplete density for the second unit. (**E**) A single modification is attached to the side chain Nδ2 atom of Asn388.

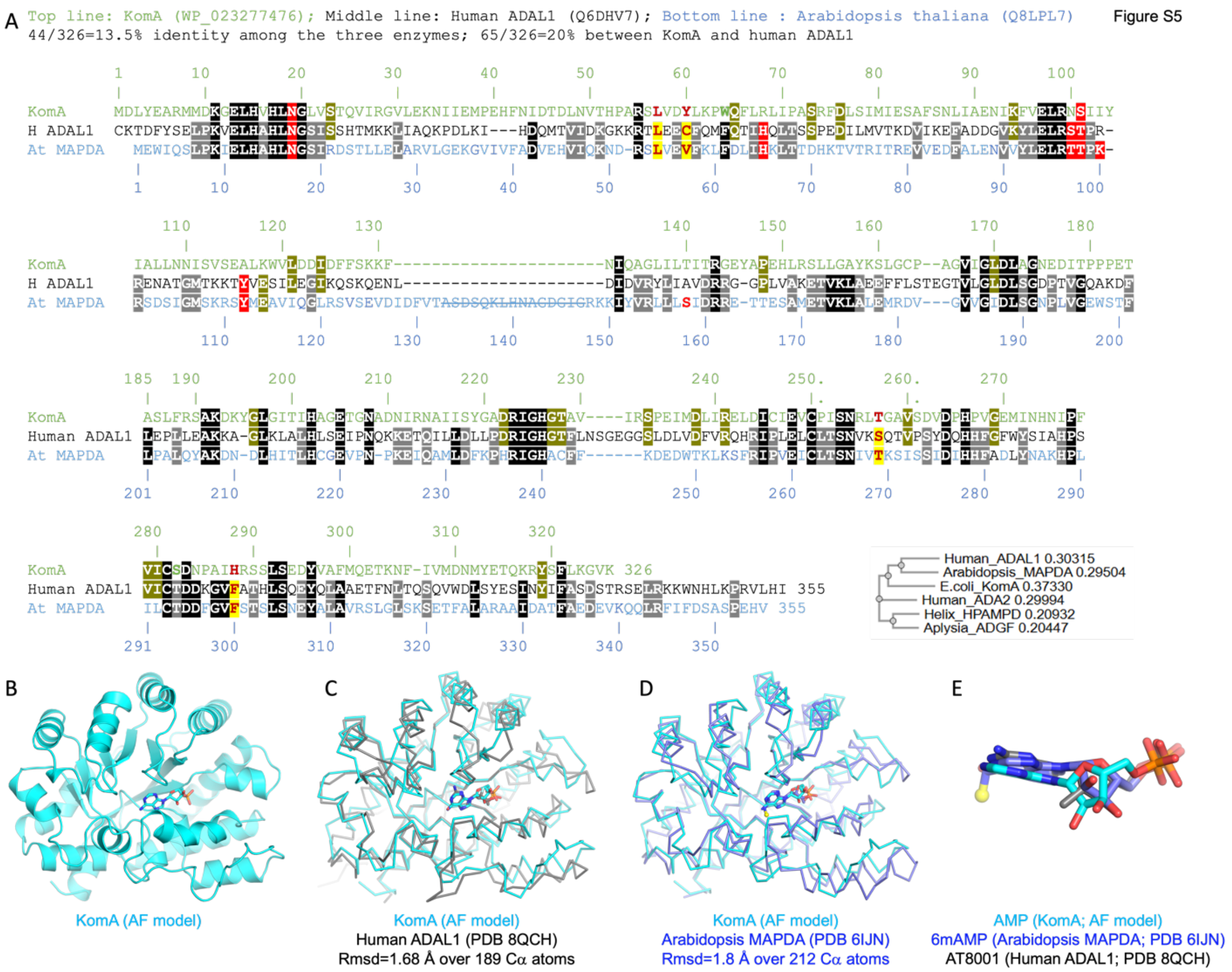


**Figure S5**. Bacterial KomA shares structural similarity to human ADAL1 and *Arabidopsis* MAPDA. (**A**) Sequence alignment of bacterial KomA (NCBI: WP_023277476.1) with two 6mAMP deaminases: human ADAL1 (UniProt: Q6DHV7) and *Arabidopsis* MAPDA (UniProt: Q8LPL7). The three enzymes have comparable lengths. Conserved residues among all three enzymes are highlighted as white letters on a black background, while those conserved between any two enzymes are shown as white letters on a grey background. Residues involved in 5'-monophosphate binding are marked as white letters on a red background, while residues that interact with the N6-methly group of 6mAMP in *Arabidopsis* MAPDA (see Figure 5B) are highlighted with a yellow background. Crossed-out residues (133-147) in *Arabidopsis* MAPDA correspond to a disordered region in the structure and are absent in KomA and human ADAL1.

Included is the phylogenetic tree of the six deaminases examined. (**B**) AlphaFold-predicted structure of KomA, with a higher confidence (pLDDT values between 70 and 90 in a scale of 100). (**C-D**) Superimposition of KomA with the experimentally determined structures of human ADAL1 and *Arabidopsis* MAPDA. (**E**) Superimposition of bound ligands. AT8001 is a guanosine analog.

Table S1. Summary of X-ray diffraction and refinement statistics of various structures.

| Deaminase | HPAMPD | | | | Chimera-2 | | |
|---|---|---|---|---|---|---|---|
| Inhibitor | - | - | Pentostatin | PMP | - | Pentostatin | PMP |
| PDB Code | 9NTE | 9NTF | 9NTG | 9NTH | 9NTI | 9NTJ | 9NTK |
| Date Collected | 06/01/2023 | 07/07/2023 | 06/01/2023 | 07/07/2023 | 06/08/2022 | 09/20/2022 | 09/23/2022 |
| Wavelength (Å) | 0.9201 | 0.9201 | 0.9201 | 0.9201 | 1.00000 | 1.00000 | 0.9201 |
| Beamline | NSLS-II Beamline 17-ID-1 | | | | APS-22ID | | NSLS-II 17-ID-1 |
| Space group | $P2_1$ | $P2_1$ | $P2_1$ | $P2_1$ | $P2_12_12_1$ | $P2_12_12_1$ | $P2_1$ |
| Unit cell (Å) | 76.28, 82.47, 211.54 | 76.34, 82.35, 211.50 | 76.12, 81.98, 212.00 | 75.99, 81.93, 212.20 | 69.45, 109.27, 173.28 | 67.35, 109.61, 171.56 | 92.83, 118.48, 109.56 |
| α, β, γ (°) | 90, 92.21, 90 | 90, 92.29, 90 | 90, 92.27, 90 | 90, 92.13, 90 | 90, 90, 90 | 90, 90, 90 | 90, 98, 90 |
| Resolution (Å) | 62.98-1.71 | 70.85-1.56 | 52.8-1.54 | 42.73-1.61 | 46.21 - 3.00 | 47.69 - 3.28 | 29.63 - 2.53 |
| [a] $R_{merge}$ | 0.143 (0.680) | 0.111 (0.800) | 0.118 (0.699) | 0.140 (0.330) | 0.454 (0.821) | 0.370 (0.550) | 0.205 (0.751) |
| $R_{pim}$ | 0.070 (0.992) | 0.065 (1.013) | 0.064 (1.003) | 0.084 (0.647) | 0.121 (0.630) | 0.123 (0.622) | 0.163 (0.985) |
| $CC_{1/2}$ | 0.994 (0.355) | 0.995 (0.320) | 0.996 (0.324) | 0.987 (0.334) | 0.985 (0.513) | 0.803 (0.331) | 0.960 (0.372) |
| [b] <I/σI> | 6.0 (0.7) | 6.0 (0.7) | 6.6 (0.6) | 5.0 (1.0) | 7.0 (1.0) | 3.1 (1.1) | 4.6 (1.4) |
| Completeness (%) | 99.9 (99.9) | 99.8 (100) | 100 (91) | 100 (98.9) | 99.3 (94.9) | 93.4 (94.9) | 97.4 (95.9) |
| Redundancy | 4.4 (3.8) | 3.8 (3.9) | 4.2 (3.5) | 3.9 (3.9) | 19.2 (9.9) | 3.9 (3.6) | 2.5 (2.5) |
| Obser. reflections | 1,249,096 | 1,430,770 | 1,480,223 | 1,296,535 | 519,122 | 73,785 | 191,520 |
| Unique reflections | 284,243 | 372,245 | 349,670 | 332,916 | 27,051 | 18,753 | 76,443 |
| **Refinement** | | | | | | | |
| Resolution (Å) | 1.71 | 1.56 | 1.54 | 1.61 | 3.00 | 3.28 | 2.53 |
| No. reflections | 283,834 | 371,300 | 349,029 | 331,929 | 26,941 | 18,669 | 76,356 |
| [c] $R_{work}$ / [d] $R_{free}$ | 0.174 / 0.202 | 0.162 / 0.188 | 0.174 / 0.196 | 0.176 / 0.202 | 0.217 / 0.275 | 0.197 / 0.273 | 0.209 / 0.260 |
| No. Atoms | | | | | | | |
| Protein | 17,112 | 16,417 | 16,465 | 16,914 | 8,023 | 8,134 | 16,324 |
| Inhibitor | - | 45[#] | 76 | 92 | - | 38 | 92 |
| Solvent | 2,100 | 2,163 | 2,193 | 2,306 | 3 | 82 | 194 |
| Zinc | 4 | 4 | 4 | 4 | 2 | 2 | 4 |
| B Factors ($Å^2$) | | | | | | | |
| Protein | 26.67 | 25.20 | 23.21 | 24.07 | 69.80 | 21.60 | 37.50 |
| Inhibitor | - | (32.00[#]) | 22.39 | 20.64 | - | 18.27 | 38.46 |
| Solvent | 40.64 | 36.70 | 33.21 | 33.92 | 54.60 | 8.09 | 30.10 |
| Zinc | 21.07 | 19.40 | 19.48 | 18.40 | 89.11 | 43.17 | 42.50 |
| **R.m.s. deviations** | | | | | | | |
| Bond lengths (Å) | 0.006 | 0.012 | 0.006 | 0.006 | 0.009 | 0.009 | 0.009 |
| Bond angles (˚) | 0.86 | 1.1 | 0.93 | 0.84 | 1.14 | 1.09 | 0.95 |
| Crystallization conditions | 0.1 M BIS-TRIS pH 6.5, 20% (w/v) PEG monomethyl ether 5,000 | 0.1 M BIS-TRIS pH 6.5, 28% (w/v) PEG monomethyl ether 2,000 | 0.1 M BIS-TRIS pH 6.5, 20% (w/v) PEG monomethyl ether 5,000 | 0.1 M BIS-TRIS pH 6.5, 20% (w/v) PEG monomethyl ether 5,000 | 0.1 M Sodium acetate trihydrate pH 4.5, 25% (w/v) PEG 3,350 | 25% PEG 1500, 15% glycerol | 0.1 M BIS-TRIS pH 5.5, 0.2 M Magnesium chloride hexahydrate, 25% (w/v) PEG 3,350 |

* Values in parenthesis correspond to highest resolution shell.

[a] $R_{merge} = \Sigma \mid I - \langle I \rangle \mid / \Sigma I$, where I is the observed intensity and <I> is the averaged intensity from multiple observations.

[b] <I/σI> = averaged ratio of the intensity (I) to the error of the intensity (σI).

[c] $R_{work} = \Sigma \mid Fobs - Fcal \mid / \Sigma \mid Fobs \mid$, where Fobs and Fcal are the observed and calculated structure factors, respectively.

[d] $R_{free}$ was calculated using a randomly chosen subset (5%) of the reflections not used in refinement.

[#] Unknown density near the active site.

**Table S2**. Primers used in the study

TRA: 5’GAT CCT GGC GGA AGC CGT ATC ATG CAA ATG CAA CGC CCC T
EFA: 5’AAG TGA GGA GTT CGT TAT CAG GCT TTT TTG AGC TAG ACG GCA
TFA: 5’ACG GCT TCC GCC AGG ATC GAC TGG GTG GTT AAA AAT GCC
ERA: 5’ ACG AAC TCC TCA CTT CGA CAT ATT GAA CAT TAT CCG CAC GAA ACT C
TFH: 5’ ACG GCT TCC GCC AGG ATC GAC TGG A
ERH: 5’ ACG AAC TCC TCA CTT CGA CAT ACT GGA CAT TG

Sequence of plasmid PD912-GAP carrying Aplysia ADGF.

```
GCTCATTCCAATTCCTTCTATTAGGCTACTAACACCATGACTTTATTAGCCTGTCTATCCTGGC
CCCCCTGGCGAGGTTCATGTTTGTTTATTTCCGAATGCAACAAGCTCCGCATTACACCCGAACA
TCACTCCAGATGAGGGCTTTCTGAGTGTGGGGTCAAATAGTTTCATGTTCCCCAAATGGCCCAA
AACTGACAGTTTAAACGCTGTCTTGGAACCTAATATGACAAAAGCGTGATCTCATCCAAGATGA
ACTAAGGATCCTTTTTTGTAGAAATGTCTTGGTGTCCTCGTCCAATCAGGTAGCCATCTCTGAA
ATATCTGGCTCCGTTGCAACTCCGAACGACCTGCTGGCAACGTAAAATTCTCCGGGGTAAAACT
TAAATGTGGAGTAATGGAACCAGAAACGTCTCTTCCCTTCTCTCTCCTTCCACCGCCCGTTACC
GTCCCTAGGAAATTTTACTCTGCTGGAGAGCTTCTTCTACGGCCCCCTTGCAGCAATGCTCTTC
CCAGCATTACGTTGCGGGTAAAACGGAGGTCGTGTACCCGACCTAGCAGCCCAGGGATGGAAAA
GTCCCGGCCGTCGCTGGCAATAATAGCGGGCGGACGCATGTCATGAGATTATTGGAAACCACCA
GAATCGAATATAAAAGGCGAACACCTTTCCCAATTTTGGTTTCTCCTGACCCAAAGACTTTAAA
TTTAATTTATTTGTCCCTATTTCAATCAATTGAACAACTATTTCCGAAACGATGAGATTCCCAT
CTATTTTCACCGCTGTCTTGTTCGCTGCCTCCTCTGCATTGGCTGCCCCTGTTAACACTACCAC
TGAAGACGAGACTGCTCAAATTCCAGCTGAAGCAGTTATCGGTTACTCTGACCTTGAGGGTGAT
TTCGACGTCGCTGTTTTGCCTTTCTCTAACTCCACTAACAACGGTTTGTTGTTCATTAACACCA
CTATCGCTTCCATTGCTGCTAAGGAAGAGGGTGTCTCTCTCGAGAAAAGAGAGGCCGAAGCTGC
TCCACTAACCAGCAAAGCCGCGTATTTACTAAAACGAAATTCTCTCATCGAGGAGGACGCCAGT
CGCAAGCTAGGCGCTAAAATTGTGCTTACTAATGAAGAAAAAGTTTTGGACGATTTCATCTTAG
CAGAAAAGCGCAAGCTGATAGATGACTCCCGCCTAAATCAAACGGAGTACATGCCTGCGGCTAG
TTTCTATCGCTCCAAAGATTTTATTGATACCACGTTTGCATACAAAATAATCCAGGACATGCCA
AAAGGAGGGGCGTTGCATTTGCATGATCTAGCAATTGCGAGTTTAGACTGGGTGGTTAAAAATG
CCACGTATAGAGACAACGTTTATATGTGCATGGACAAAGACAATGATGTCAATCTACGAGTTCT
GCAGCTGATCCCGCCAGATCCTTTCTGTGTGTGGAAATTAGTTGCCACAGAACGCGCCAACTCA
GGAGACGTCGAAGCATTTGACGATTGGCTCAAAAAGAACATCTCGTATCTCTCTACGGATCCTG
TCACGCAGTATGCCACTGTTGATTCAGTCTGGGTCAGGTTCAACAAGTACTTCGCCCAAGTCAT
AGGGCTGCTCTTCTACGCGCCGATCATGAGAGATTACTACCGTCAAGCTCTGGAAGAGTTTCGT
GCGGATAATGTTCAATATATTGAACTGAGATCGCAGCTCTTCGGCTTTTTTGAGCTAGACGGCA
CAGTTCATGACGCGGAATTTGGTCTTAATCTCTACAAGTCTGTAACGGAAGAGTTCCAGCGAGA
ATATCCCGATTTTATCGGTGCGAAGATTATTCTGTCTGGTTTACGATTCAAGTCTCAGGAGGAA
ATTTTGAACGAGGTAAAAATCGCCATGGATCTGCACAAGAAATACCCAGACTTCTTCCTGGGAT
ATGATCTCGTAGGCCAAGAAGATCCGAATTTTTCCCTGCTTCATTATTTGGATGCACTGCTGTA
TCCGTCAATACAAAATCCGCCATATCGGCTACCGTATTTTTTCCACGCTGCAGAGACCAACTGG
CAGGAAACTGAAGTCGACTACAACCTTGCAGACGCACTCCTGCTCAATACCACTCGAGTCGGCC
ATGGGTTTGCATTAATCAAGCATCCCCGGTTCACAGAGCTTGCAAAGGAAAACGGTGTGGCCGT
GGAAGTGAATCCAATTTCCAACCAGATTCTTGGGTTGGTAAGGGATGTTCGCAACCACGCTTTG
GTGCCTCTGATTGCCGACGACTATCCCATTGTGATATCGAGCGACGACCCTGGAGCTTGGGAGG
CTTCTCCCCTGTCACACGACTTTTATGTAGCGCTGATGGACCTGTGTGGTCGGGATACGGCATT
```

AACATTTTTGAAACAACTCGCCTTAAATTCAATTAGATATTCCGCAATGAGCGATACCGAGAAG
GTCGCTGCAAAGGCAAAATGGACTACACAATGGGACAAGTTTGTCAAGACGTCGGTGGAGGGGT
TAAAGCCACATATAAACGACAGATCACACCACCACCACCACCACCACCACCACCACTGAGGTGA
GGTTGAAGGGGCGGCCGCTCAAGAGGATGTCAGAATGCCATTTGCCTGAGAGATGCAGGCTTCA
TTTTTGATACTTTTTTATTTGTAACCTATATAGTATAGGATTTTTTTTGTCATTTTGTTTCTTC
TCGTACGAGCTTGCTCCTGATCAGCCTATCTCGCAGCAGATGAATATCTTGTGGTAGGGGTTTG
GGAAAATCATTCGAGTTTGATGTTTTTCTTGGTATTTCCCACTCCTCTTCAGAGTACAGAAGAT
TAAGTGAAACCTTCGTTTGTGCGGATCCTTCAGTAATGTCTTGTTTCTTTTGTTGCAGTGGTGA
GCCATTTTGACTTCGTGAAAGTTTCTTTAGAATAGTTGTTTCCAGAGGCCAAACATTCCACCCG
TAGTAAAGTGCAAGCGTAGGAAGACCAAGACTGGCATAAATCAGGTATAAGTGTCGAGCACTGG
CAGGTGATCTTCTGAAAGTTTCTACTAGCAGATAAGATCCAGTAGTCATGCATATGGCAACAAT
GTACCGTGTGGATCTAAGAACGCGTCCTACTAACCTTCGCATTCGTTGGTCCAGTTTGTTGTTA
TCGATCAACGTGACAAGGTTGTCGATTCCGCGTAAGCATGCATACCCAAGGACGCCTGTTGCAA
TTCCAAGTGAGCCAGTTCCAACAATCTTTGTAATATTAGAGCACTTCATTGTGTTGCGCTTGAA
AGTAAAATGCGAACAAATTAAGAGATAATCTCGAAACCGCGACTTCAAACGCCAATATGATGTG
CGGCACACAATAAGCGTTCATATCCGCTGGGTGACTTTCTCGCTTTAAAAAATTATCCGAAAAA
ATTTTCTAGAGTGTTGTTACTTTATACTTCCGGCTCGTATAATACGACAAGGTGTAAGGAGGAC
TAAACCATGGCTAAACTCACCTCTGCTGTTCCAGTCCTGACTGCTCGTGATGTTGCTGGTGCTG
TTGAGTTCTGGACTGATAGACTCGGTTTCTCCCGTGACTTCGTAGAGGACGACTTTGCCGGTGT
TGTACGTGACGACGTTACCCTGTTCATCTCCGCAGTTCAGGACCAGGTTGTGCCAGACAACACT
CTGGCATGGGTATGGGTTCGTGGTCTGGACGAACTGTACGCTGAGTGGTCTGAGGTCGTGTCTA
CCAACTTCCGTGATGCATCTGGTCCAGCTATGACCGAGATCGGTGAACAGCCCTGGGGTCGTGA
GTTTGCACTGCGTGATCCAGCTGGTAACTGCGTGCATTTCGTCGCAGAAGAACAGGACTAACAA
TTGACACCTTACGATTATTTAGAGAGTATTTATTAGTTTTATTGTATGTATACGGATGTTTTAT
TATCTATTTATGCCCTTATATTCTGTAACTATCCAAAAGTCCTATCTTATCAAGCCAGCAATCT
ATGTCCGCGAACGTCAACTAAAAATAAGCTTTTTATGCTGTTCTCTCTTTTTTTCCCTTCGGTA
TAATTATACCTTGCATCCACAGATTCTCCTGCCAAATTTTGCATAATCCTTTACAACATGGCTA
TATGGGAGCACTTAGCGCCCTCCAAAACCCATATTGCCTACGCATGTATAGGTGTTTTTTCCAC
AATATTTTCTCTGTGCTCTCTTTTTATTAAAGAGAAGCTCTATATCGGAGAAGCTTCTGTGGCC
GTTATATTCGGCCTTATCGTGGGACCACATTGCCTGAATTGGTTTGCCCCGGAAGATTGGGGAA
ACTTGGATCTGATTACCTTAGCTGCAGGTACCACTGAGCGTCAGACCCCGTAGAAAAGATCAAA
GGATCTTCTTGAGATCCTTTTTTTCTGCGCGTAATCTGCTGCTTGCAAACAAAAAAACCACCGC
TACCAGCGGTGGTTTGTTTGCCGGATCAAGAGCTACCAACTCTTTTTCCGAAGGTAACTGGCTT
CAGCAGAGCGCAGATACCAAATACTGTTCTTCTAGTGTAGCCGTAGTTAGGCCACCACTTCAAG
AACTCTGTAGCACCGCCTACATACCTCGCTCTGCTAATCCTGTTACCAGTGGCTGCTGCCAGTG
GCGATAAGTCGTGTCTTACCGGGTTGGACTCAAGACGATAGTTACCGGATAAGGCGCAGCGGTC
GGGCTGAACGGGGGGTTCGTGCACACAGCCCAGCTTGGAGCGAACGACCTACACCGAACTGAGA
TACCTACAGCGTGAGCTATGAGAAAGCGCCACGCTTCCCGAAGGGAGAAAGGCGGACAGGTATC
CGGTAAGCGGCAGGGTCGGAACAGGAGAGCGCACGAGGGAGCTTCCAGGGGGAAACGCCTGGTA
TCTTTATAGTCCTGTCGGGTTTCGCCACCTCTGACTTGAGCGTCGATTTTTGTGATGCTCGTCA
GGGGGGCGGAGCCTATGGAAAAACGCCAGCAACGCGGCCTTTTTACGGTTCCTGGCCTTTTGCT
GGCCTTTTGCTCACATGTTCTTTCCTGCGGTACCCAGATCCAATTCCCGCTTTGACTGCCTGAA
ATCTCCATCGCCTACAATGATGACATTTGGATTTGGTTGACTCATGTTGGTATTGTGAAATAGA
CGCAGATCGGGAACACTGAAAAATACACAGTTATTATTCATTTAAATAACATCCAAAGACGAAA
GGTTGAATGAAACCTTTTTGCCATCCGACATCCACAGGTCCATTCTCACACATAAGTGCCAAAC
GCAACAGGAGGGGATACACTAGCAGCAGACCGTTGCAAACGCAGGACCTCCACTCCTCTTCTCC
TCAACACCCACTTTTGCCATCGAAAAACCAGCCCAGTTATTGGGCTTGATTGGAGCTC
